# Rosace: a robust deep mutational scanning analysis framework employing position and mean-variance shrinkage

**DOI:** 10.1101/2023.10.24.562292

**Authors:** Jingyou Rao, Ruiqi Xin, Christian Macdonald, Matthew Howard, Gabriella O. Estevam, Sook Wah Yee, Mingsen Wang, James S. Fraser, Willow Coyote-Maestas, Harold Pimentel

**Affiliations:** Department of Computer Science, UCLA, Los Angeles, CA, USA; Computational and Systems Biology Interdepartmental Program, UCLA, Los Angeles, CA, USA; Department of Bioengineering and Therapeutic Sciences, UCSF, San Francisco, CA, USA; Tetrad Graduate Program, UCSF, San Francisco, CA, USA; Department of Pharmaceutical Chemistry, UCSF, San Francisco, CA, USA; Department of Mathematics, Baruch College, CUNY, New York, NY, USA; Quantitative Biosciences Institute, UCSF, San Francisco, CA, USA; Department of Computational Medicine, David Geffen School of Medicine, UCLA, Los Angeles, CA, USA; Department of Human Genetics, David Geffen School of Medicine, UCLA, Los Angeles, CA, USA

## Abstract

Deep mutational scanning (DMS) enables functional insight into protein mutations with multiplexed measurements of thousands of genetic variants in a protein simultaneously. The small sample size of DMS renders classical statistical methods ineffective, for example, p-values cannot be correctly calibrated when treating variants independently. We propose `Rosace`, a Bayesian framework for analyzing growth-based deep mutational scanning data. `Rosace` leverages amino acid position information to increase power and control the false discovery rate by sharing information across parameters via shrinkage. To benchmark `Rosace` against existing methods, we developed `Rosette`, a simulation framework that simulates the distributional properties of DMS. Further, we show that `Rosace` is robust to the violation of model assumptions and is more powerful than existing tools under `Rosette` simulation and real data.

## 1 Background

Understanding how protein function is encoded at the residue level is a central challenge in modern protein science. Mutations can cause diseases and drive evolution through perturbing protein function in a myriad of ways, such as by altering its conformational ensemble and stability or its interaction with ligands and binding partners. In these contexts, mutations may result in a loss of function, gain of function, or a neutral phenotype (i.e., no discernable effects). Mutations also often exert effects across multiple phenotypes, and these perturbations can ultimately propagate to alter complex processes in cell biology and physiology. Reverse genetics approaches offer a powerful handle for researchers to investigate biology via introducing mutations and observing the resulting phenotypic changes.

Deep mutational scanning (DMS) is a technique for systematically determining the effect of a large library of mutations individually on a phenotype of interest by performing pooled assays and measuring the relative effects of each variant (Fig. 1A). Taking enzymes as an example, these phenotypes could include catalytic activity [1] or stability [2]. For a transcription factor, the phenotype could be DNA binding specificity or transcriptional activity [3]. The relevant phenotype for a membrane transporter might be folding and trafficking or substrate transport [4]. These phenotypes are often captured by either growth-based [2, 4, 5, 6, 7, 8] or fluorescence-based assays [3, 4, 9]. Those two experiments are inherently differently designed and merit separate analysis frameworks. In growth-based assays, the relative growth rates of cells are of interest. In fluorescence-based assays, changes to the distribution of reporter gene expression are measured. In this paper, we focus solely on growth-based screens.

**Figure 1.**
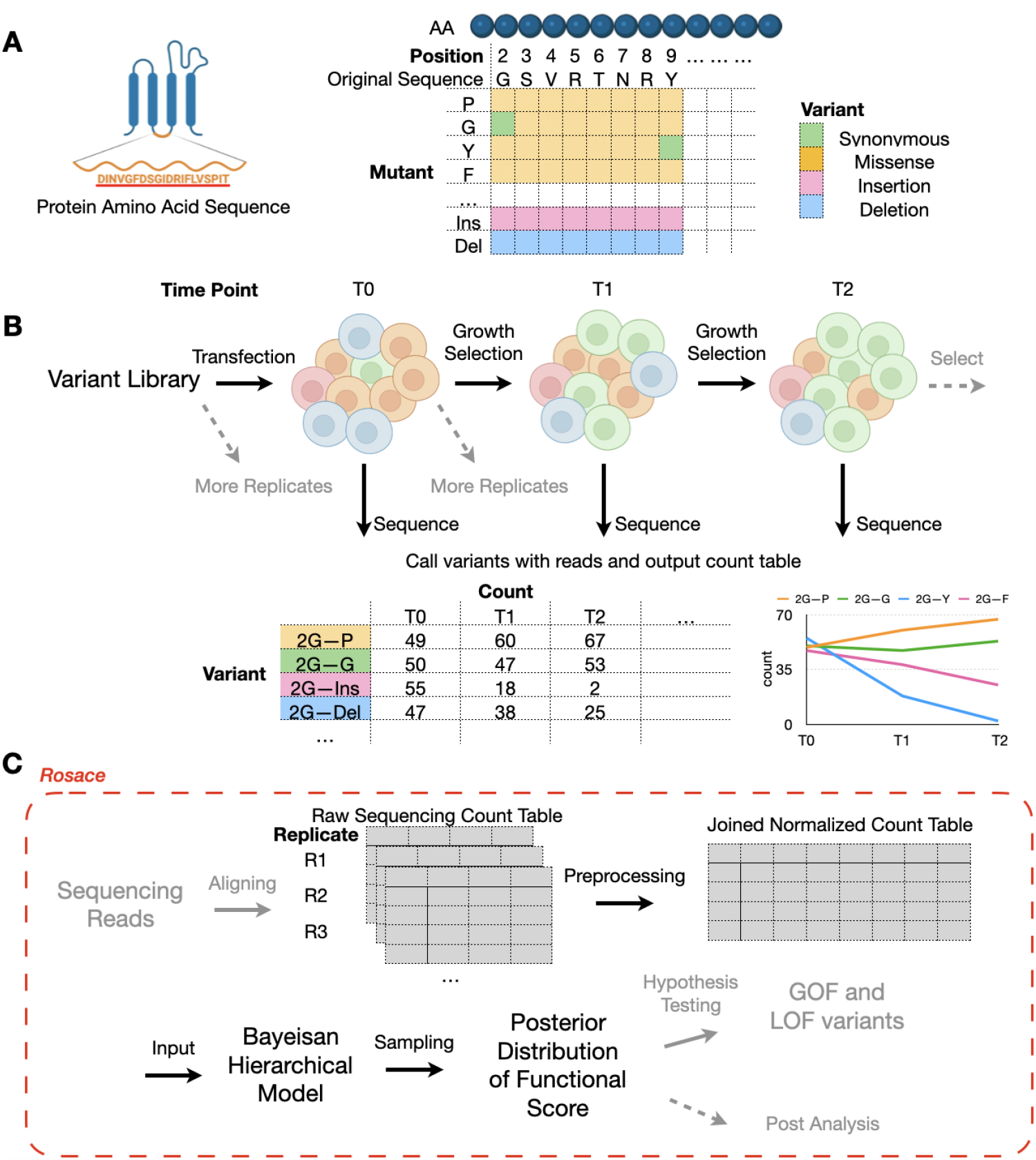
Deep Mutational Scanning and Overview of `Rosace` Framework. (A) Each amino acid of the selected protein sequence is mutated to another mutant in deep mutational scanning. (B) Cells carrying different variants are grown in the same pool under selection pressure. At each time point, cells are sequenced to output the count table. (C) `Rosace` is an R package that accepts input from the raw sequencing count table and outputs the posterior distribution of functional score.

In a growth-based DMS experiment, we grow a pool of cells carrying different variants under a selective pressure linked to gene function. At set intervals, we sequence the cells to identify each variant’s frequency in the pool. The change in the frequency over the course of the experiment, from initial frequencies to subsequent measurements serves as a metric of the variant’s functional effects (Fig. 1B). The functional score is often computed for each variant in the DMS screen and compared against those of synonymous mutations or wild-type cells to display the relative functional change of the protein caused by the mutation. Thus, reliable inference of functional scores is crucial to understanding both the individual mutation as well as at which residue location variants tend to have significant functional effects.

The main challenge of functional score inference is that even under the simplest model, there are at least two estimators required for each mutation (mean and variance of functional change), and in practice, it is rare to have more than three replicates. As a result, it has been posited that under naïve estimators that have been commonly employed, there are likely issues with the false discovery rate and the statistical power of detecting mutations that significantly change the function of the protein [10]. Regardless, incorporating domain-specific assumptions is required to make inference tractable with few samples and thousands of parameters.

To alleviate the small-sample-size inference problem in DMS [11], three commonly used methods have been developed: *Enrich2* [12], *DiMSum* [10], and *EMPIRIC* [13]. *Enrich2* simplifies the variance estimator by assuming that counts are Poisson-distributed (the variance being equal to the mean) and combines the replicates using a random-effect model. *DiMSum*, however, argues that the assumption in *Enrich2* is not enough to control type-I error. As a result, *DiMSum* builds upon *Enrich2* and includes additional variance terms to model the over-dispersion of sequencing counts. However, as presented in Faure et al. 2020 [10], this ratio-based method only applies to the DMS screen with one round of selection, while more recent DMS screens might have more than two rounds of selection (i.e. sampling at multiple time points) [4, 5]. Alternatively, *EMPIRIC* fits a Bayesian model that infers each variant separately with non-informative uniform priors to all parameters and thus does not shrink the estimates to robustly correct the variance in estimates due to the small sample size. Further, the model does not accommodate multiple replicates. In addition, *mutscan* [14], a recently developed R package for DMS analysis, employed two established statistical models *edgeR* and *limma-voom*. However, these two methods were originally designed for RNA-seq data and the data generation process for DMS is very different. One of the key differences is consistency among replicates. In RNA-seq, gene expression is relatively consistent across replicates under the same condition, while in DMS, counts of variants can vary much since the a priori representation in the initial variant library can be vastly inconsistent among replicates.

While these methods provide reasonable regularization of the score’s variance, additional information can further improve the prior. One solution is incorporating residue position information. It has been noted that amino acids in particular regions have an oversized effect on the protein’s function, and thus analyzing DMS in a position-specific manner has resulted in several interesting findings [15, 16]. In the form of hidden Markov models (HMMs) and position-specific scoring matrices (PSSMs), this is the basis for the sensitive detection of homology in protein sequences [17]. These results directly imply that variants at the same position likely share some similarities in their behavior and thus that incorporating local information into modeling might produce more robust inferences. However, no existing methods have incorporated residue position information into their models yet.

To overcome these limitations, we present `Rosace`, the first growth-based DMS method that incorporates local positional information to increase inference performance. `Rosace` implements a hierarchical model that parameterizes each variant’s effect as a function of the positional effect, thus providing a way to incorporate both position-specific information and shrinkage into the model. Additionally, we developed `Rosette`, a simulation framework that attempts to simulate several properties of DMS such as bimodality, similarities in behavior across similar substitutions, and the overdispersion of counts. We use `Rosette` to simulate several screening modalities and show that our inference method, `Rosace`, exhibits higher power and controls the false discovery rate (FDR) better on average than existing methods. Importantly, `Rosace` and `Rosette` are not two views of the same model - `Rosette` is based on a set of assumptions that are different from or even opposite to those of `Rosace`. `Rosace`’s ability to accommodate data generated under different assumptions shows its robustness. Finally, we run `Rosace` on real datasets and it shows a much lower FDR than existing methods while maintaining similar power on experimentally validated positive controls.

## 2 Results

### 2.1 Overview of Rosace Framework

`Rosace` is a Bayesian framework for analyzing growth-based deep mutational scanning data, producing variant-level estimates from sequencing counts. The method requires as input the raw sequencing counts. It outputs the posterior distribution of variants’ functional scores, which can be further evaluated to conduct hypothesis testing, plotting, and other downstream analyses (Fig. 1C). `Rosace` is available as an R package. To generate the input of `Rosace` from sequencing reads, we share a Snakemake workflow dubbed Dumpling for short-read-based experiments in the GitHub repository described in the Methods section. Additionally, `Rosace` supports input count data processed from *Enrich2* [12] for other protocols.

### 2.2 Rosace Hierarchical Model with Positional Information and Score Shrinkage

Here, we begin by motivating the use of positional information. Next, we describe the intuition of how we use the positional information. Finally, we describe the remaining dimensions of shrinkage which assist in robust estimates with few experiment replicates.

A variant is herein defined as the amino acid identity at a position in a protein, where that identity differs from the wild-type sequence. The sequence position of a variant *p*(*v*) provides information on the functional effects to the protein from the variant. We define the position-level functional score *ϕ*_*p*(*v*)_ as the mean functional score of all variants on a given position.

To motivate the use of positional information, we take the posterior distribution of the position-level functional score estimated from a real DMS experiment, a cytotoxicity-based growth screen of a human transporter, OCT1 (Fig. 2A). In this experiment, variants with decreased activity are expected to increase in abundance, as they lose the ability to import a cytotoxic substrate during selection, and variants with increased activity will decrease in abundance similarly. We observe that most position-level score estimates 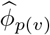 significantly deviate from the mean, implying that position has material idiosyncratic variation and thus carries information about the protein’s functional architecture.

**Figure 2.**
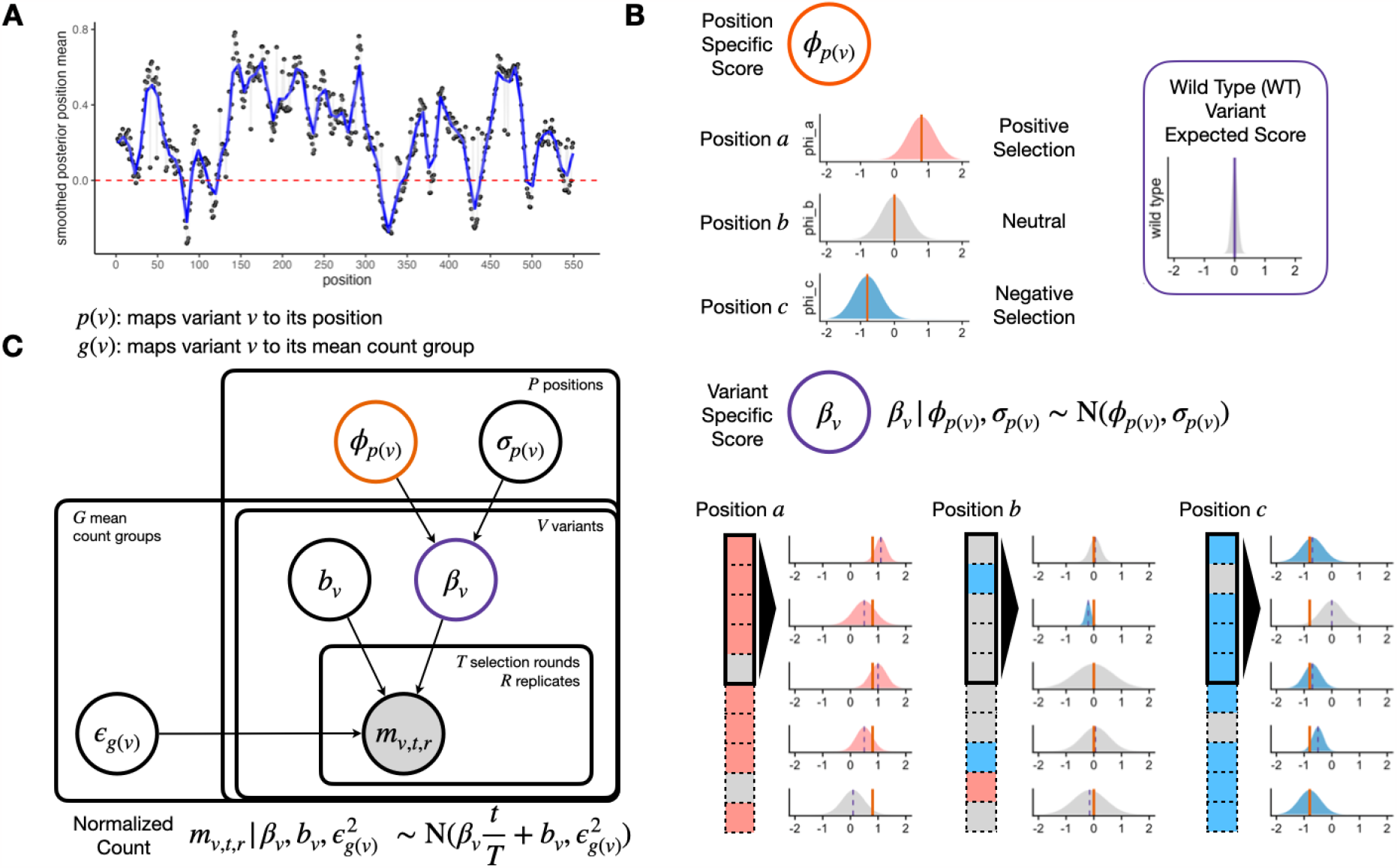
`Rosace` shares information at the same position to inform variant effects. (A) Smoothed position-specific score (sliding window = 5) across positions from OCT1 cytotoxicity screen. Red dotted lines at score = 0 (neutral position). (B) A conceptual view of the `Rosace` generative model. Each position has an overall effect, from which variant effects are conferred. Note the prior is wide enough to allow effects that do not follow the mean. Wild type score distribution is assumed to be at 0. (C) Plate model representation of `Rosace`. See Methods for the description of parameters.

To incorporate the positional information into our model, we introduce a position-specific score *ϕ*_*p*(*v*)_ where *p*(*v*) maps variant *v* to its amino acid position. The variant-specific score *β*_*v*_ is regularized and controlled by the value of *ϕ*_*p*(*v*)_. To illustrate the point, we conceptually categorize position into three types: positively selected (*ϕ*_*p*(*v*)_ ≫ 0), (nearly) neutral (*ϕ*_*p*(*v*)_ ≈ 0), and negatively selected (*ϕ*_*p*(*v*)_ ≪ 0) (Fig. 2B). Variants in a positively selected position tend to have scores centered around the positive mean estimate of *ϕ*_*p*(*v*)_, and vice versa for the negatively selected position. Variants in a neutral position tend to be statistically non-significant as the region might not be important to the measured phenotype.

Regularization of the score’s variance is achieved mainly by sharing information across variants within the position and asserting weakly informative priors on the parameters (Fig. 2C). Functional scores of the variants within the position are drawn from the same set of parameters *ϕ*_*p*(*v*)_ and *σ*_*p*(*v*)_. The error term *ϵ*_*g*(*v*)_ in the linear regression on normalized counts is also shared in the mean count group (see in Methods) to prevent biased estimation of the error and incorporate mean-variance relationship commonly modeled in RNA-seq [18, 19]. Importantly, while we use the position information to center the prior, the prior is weak enough to allow variants at a position to deviate from the mean. The variant-level intercept *b*_*v*_ is given a strong prior with a tight distribution centered at 0 to prevent over-fitting.

### 2.3 Rosace Performance on OCT1 Cytotoxicity Data

To test the performance of `Rosace`, we ran `Rosace` along with *Enrich2, mutscan* (both *limma-voom* and *edgeR*) and simple linear regression (the naïve method) on the OCT1 cytotoxicity screen. *DiMSum* is not included in this comparison because it cannot analyze data with three selection rounds. The data is pre-processed with wild-type normalization for all three methods. The analysis is done on all subsets of three replicates ({1}, {2}, {3}, {1, 2}, {1, 3}, {2, 3}, {1, 2, 3}).

While we do not have a set of true negative control variants, we assume most synonymous mutations would not change the phenotype, and thus, we use synonymous mutation as a proxy for negative controls. We compute the percentage of significant synonymous mutations called by the hypothesis testing as one representation of the false discovery rate (FDR). The variants are ranked based on the hypothesis testing statistics from the method (p-value for frequentist methods and local false sign rate [20], or *lfsr*) for Bayesian methods). In an ideal scenario with no noise, the line of ranked variants by FDR is flat at 0 and slowly rises after all true variants with effect are called. `Rosace` has a very flat segment among the top 25% of the ranked variants compared to *Enrich2* and the naive method and keeps the FDR lower than *limma-voom* and *edgeR* until the end (Fig. 3A). Importantly, we note that the `Rosace` curve moves only slightly from 1 replicate to 3 replicates, while the other methods shift more, implying that the change in the number of synonymous mutations called is minor for `Rosace`, despite having fewer replicates (Fig. 3A).

**Figure 3.**
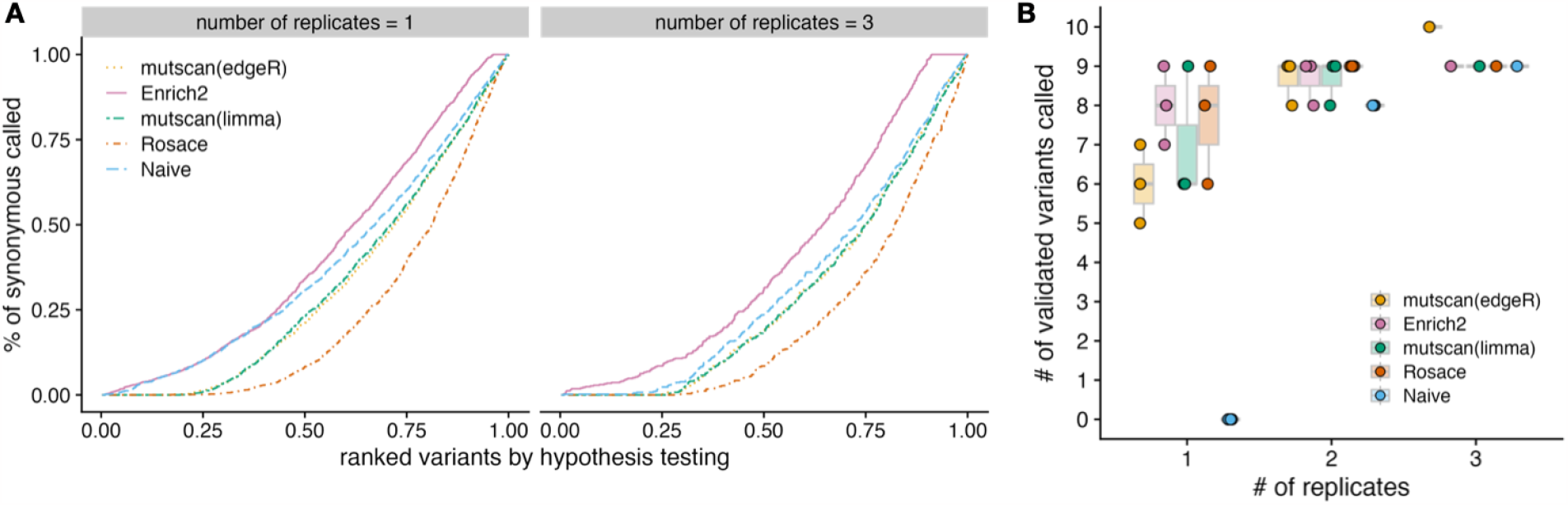
False Discovery Rate and Sensitivity on OCT1 Cytotoxicity Data. (A) Percent of synonymous mutations called (false discovery rate) versus ranked variants by hypothesis testing. The left panel is from taking the mean of analysis of the three individual replicates. Ideally, the line would be flat at 0 until all the variants with true effects are discovered. (B) Number of validated variants called (in total 10) versus number of replicates. If only 1 or 2 replicates are used, we iterate through all possible combinations. For example, the three points for `Rosace` on 2 replicates use Replicate {1, 2}, {1, 3}, and {2, 3} respectively.

While lower FDR may result in lower power in the method, we show that `Rosace` is consistently powerful in detecting the OCT1 positive control variants. Yee, Macdonald, and Mitrovic *et al*. [4] conducted lower-throughput radioligand uptake experiments in HEK293T cells and validated 10 variants that have a loss-of-function or gain-of-function phenotype. We use the number of validated variants to approximate the power of the method. As shown in Figure 3B, `Rosace` has comparable power to *Enrich2, limma-voom*, and *edgeR* regardless of the number of replicates, while the naive method is unable to detect anything in the case of one replicate. `Rosace` calls significantly fewer synonymous mutations than every other method while maintaining high power, showing that `Rosace` is robust in real data.

To test `Rosace` performance on diverse datasets, we run `Rosace` on the CARD11 data [8] (5 replicates, 1 selection round, 2.6k variants), the MSH data [6] (3 replicates, 1 selection round, 18k variants), and the BRCA1 data [7] (2 replicates, 2 selection rounds, 3.9k variants). `Rosace` consistently shows high sensitivity in detecting the positive control variants in all three datasets while controlling the false discovery rate (Fig. S1).

### 2.4 Rosette: DMS data simulation which matches marginal distributions from real DMS data

To further benchmark the performance of `Rosace` and other related methods, we propose a new simulation framework called `Rosette`, which generates DMS data using parameters directly inferred from the real experiment to gain the flexibility of mimicking the overall structure of most growth-based DMS screen data (Fig. 4A).

**Figure 4.**
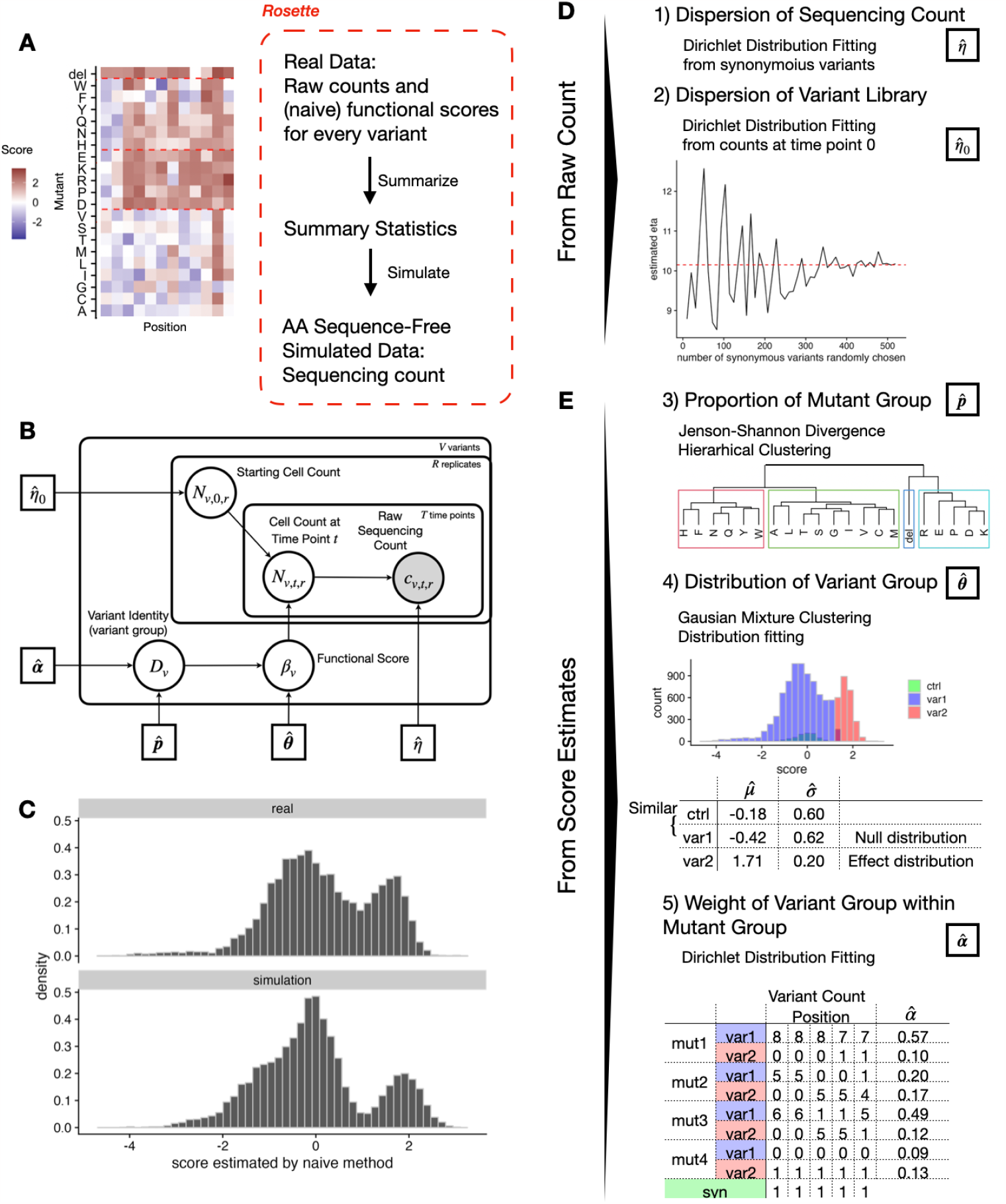
`Rosette` simulation framework preserves the overall structure of growth-based DMS screens. The plots show the result of using OCT1 data as input. (A) `Rosette` generates summary statistics from real data and simulates the sequencing count. (B) Generative model for `Rosette` simulation. (C) The distribution of real and predicted functional scores is similar. (D) (E) Five summary statistics are needed for `Rosette`.

Intuitively, if we construct a simulation that closely follows the assumptions of our model, our model should have outstanding performance. To facilitate a fair comparison with other methods, the simulation presented here is not aligned with the assumptions made in `Rosace`. In fact, the central assumption that variant position carries information is violated by construction to showcase the robustness of `Rosace`.

To re-clarify the terminology used throughout this paper, “mutant” refers to the substitution, insertion, or deletion of amino acids. A position-mutant pair is considered a variant. Mutants are categorized into mutant groups with hierarchical clustering schemes or predefined criteria (our model uses the former that are expected to align with the biophysical properties of amino acids). Variants are grouped in two ways: 1) by their functional change to the protein, namely neutral, loss-of-function (LOF), or gain-of-function (GOF), referred to as “variant groups”, and 2) by the mean of the raw sequencing counts across replicates, referred to as “variant mean groups”.

`Rosette` calculates two summary statistics from the raw sequencing counts (dispersion of the sequencing count *η* and dispersion of the variant library *η*_0_) (Fig. 4D) and three others from the score estimates (the proportion of each mutant group p, the functional score’s distribution of each variant group θ, and the weight of each variant group α) (Fig. 4E). Since we are only learning the distribution of the scores instead of the functional characteristics of individual variants, the score estimates can be naïve (e.g. simple linear regression) or more complicated (e.g. `Rosace`).

The dispersion of the sequencing counts *η* measures how much variability in variant representation there is in the entire experimental procedure, during both cell culture and sequencing. When *η* goes to infinity, it means that the sequencing count is almost the same as the expected true cell count (no overdispersion). When *η* is small, it shows an over-dispersion of the sequencing count. In an ideal experiment with no overdispersion, the proportion of synonymous mutations should be invariant to time due to the absence of functional changes. However, from the real data, we have observed a large variability of proportion changes within the synonymous mutations at different selection rounds, which is attributed to overdispersion and cannot be explained by a simple multinomial distribution (Fig. S2). Therefore, we choose to model the sequencing step with a Dirichlet-Multinomial distribution that includes *η* as the dispersion parameter.

The dispersion of variant library *η*_0_ measures how much variability already exists in variant representation before the cell selection. Theoretically, each variant would have around the same number of cells at the initial time point. However, due to the imbalance during the variant library generation process and the cell culture of the initial population that might already be under selection, we sometimes see a wide dispersion of counts across variants. To estimate this dispersion, we fit a Dirichlet-Multinomial distribution under the assumption that the proportion of synonymous variants in the cell pool should be invariant throughout the experiment.

The distribution and the structure of the underlying true functional score across variants are controlled by the rest of the summary statistics. We make a few assumptions here. First, the functional score distribution of mutants across positions (or a row in the heatmap (Fig. 4A)) is different, but within the mutant group, the mutants are independent and identically distributed (or exchangeable). We estimate the mutant group by hierarchical clustering with distance defined by empirical Jenson-Shannon Divergence and record its proportion 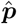. Second, each variant belongs to the neutral hypothesis (score close to 0, similar to synonymous mutations) or the alternative hypothesis (away from 0, different from synonymous mutations). The number of the variant group can be 1-3 (neutral, GOF, and LOF) based on the number of modes in the marginal functional score distribution, and the variants within a variant group are exchangeable. We estimate the borderline of the variant group by Gaussian mixture clustering and fit the distribution parameter 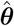. Finally, we assume that the positions are independent. While this is a simplifying assumption, to consider the relationship between positions we would need to incorporate additional assumptions about the functional region of the protein. As a result, we treat the positions as exchangeable and model the proportion of variant group identity (neutral, GOF, LOF) in each mutant group by a Dirichlet distribution with parameter 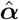.

To simulate the sequencing count from the summary statistics, we use a generative model that mimics the experiment process and is completely different from the `Rosace` inference model for fair benchmarking. We first draw the functional score of each variant *β*_*v*_ from the structure described in the summary statistics and the ones in the neutral group are set to be 0. Then, we map the functional score to its latent functional parameters: the cell growth rate in the growth screen. Next, we generate the cell count at a particular time point *N*_*v,t,r*_ by the cell count at the previous time point *N*_*v,t*−1,*r*_ and the latent functional parameters. Finally, the sequencing count is generated from a Dirichlet-Multinomial distribution with the summarized dispersion parameter and the cell count.

The simulation result shows that the simulated functional score distribution is comparable to the real experimental data (Fig. 4C). We also demonstrate that the simulation is not particularly favorable to models containing positional information such as `Rosace`. From Figure 4E, we observe that in the simulation, the positional-level score is not as widespread as the real data. In addition, the positions with extreme scores (very positive scores in the OCT1 dataset) have reduced standard deviation in the real data, but not in the simulation (Fig. S3). As a result, we would expect the performance of `Rosace` to be better in real data than in the simulation.

### 2.5 Testing Rosace False Discovery Control with Rosette Simulation

To test the performance of `Rosace`, we generate simulated data using `Rosette` from two distinctive growth-based assays: the transporter OCT1 data where LOF variants are positively selected [4] and the kinase MET data where LOF variants are negatively selected [5]. The OCT1 DMS screen measures the impact of variants on cytotoxic drug SM73 uptake mediated by the transporter OCT1. If a mutation causes the transporter protein to have decreased activity, the cells in the pool will import less substrate and thus die more slowly than wide-type or those with synonymous mutations, so the LOF variants would be positively selected. In the MET DMS screen, the kinase drives proliferation and cell growth in the BA/F3 mammalian cell line in the absence of IL-3 (interleukin-3) withdrawal. If the variant protein fails to function, the cells will die faster than the wild-type cells, so the LOF variants will be negatively selected. Both data sets have a clear separation of two modes in the functional score distribution (neutral and LOF) (Fig. S4). We benchmark `Rosace` with *Enrich2, edgeR, limma-voom*, and the naive method in scenarios where we use 1 or all 3 of replicates and 1 or all 3 of selection rounds. *DiMSum* is benchmarked when there is only one round of selection because it is not designed to handle multiple rounds. Each scenario is repeated 10 times. The results of all methods show similar correlations with the latent growth rates (Fig. S5), and thus, for benchmarking purposes, we focus on hypothesis testing.

We compare methods from a variant ranking point of view, comparing methods in terms of the number of false discoveries for any given number of variants selected to be LOF. This is because `Rosace` is a Bayesian framework that uses *lfsr* instead of p-values as the metric for variant selection and it is hard to translate *lfsr* to FDR for a hard threshold. Variants are ranked by adjusted p-values or *lfsr* (ascending). Methods that perform well will rank the truly LOF variants in the simulation ahead of non-LOF variants. In an ideal scenario with no noise, we would expect the line of ranked variants by FDR to be flat at 0 and slowly rise after all LOF variants are called. The results in Figure 5 show that even though the position assumption is violated in the `Rosette` simulation, `Rosace` is robust enough to maintain relatively low FDR in all simulation conditions.

**Figure 5.**
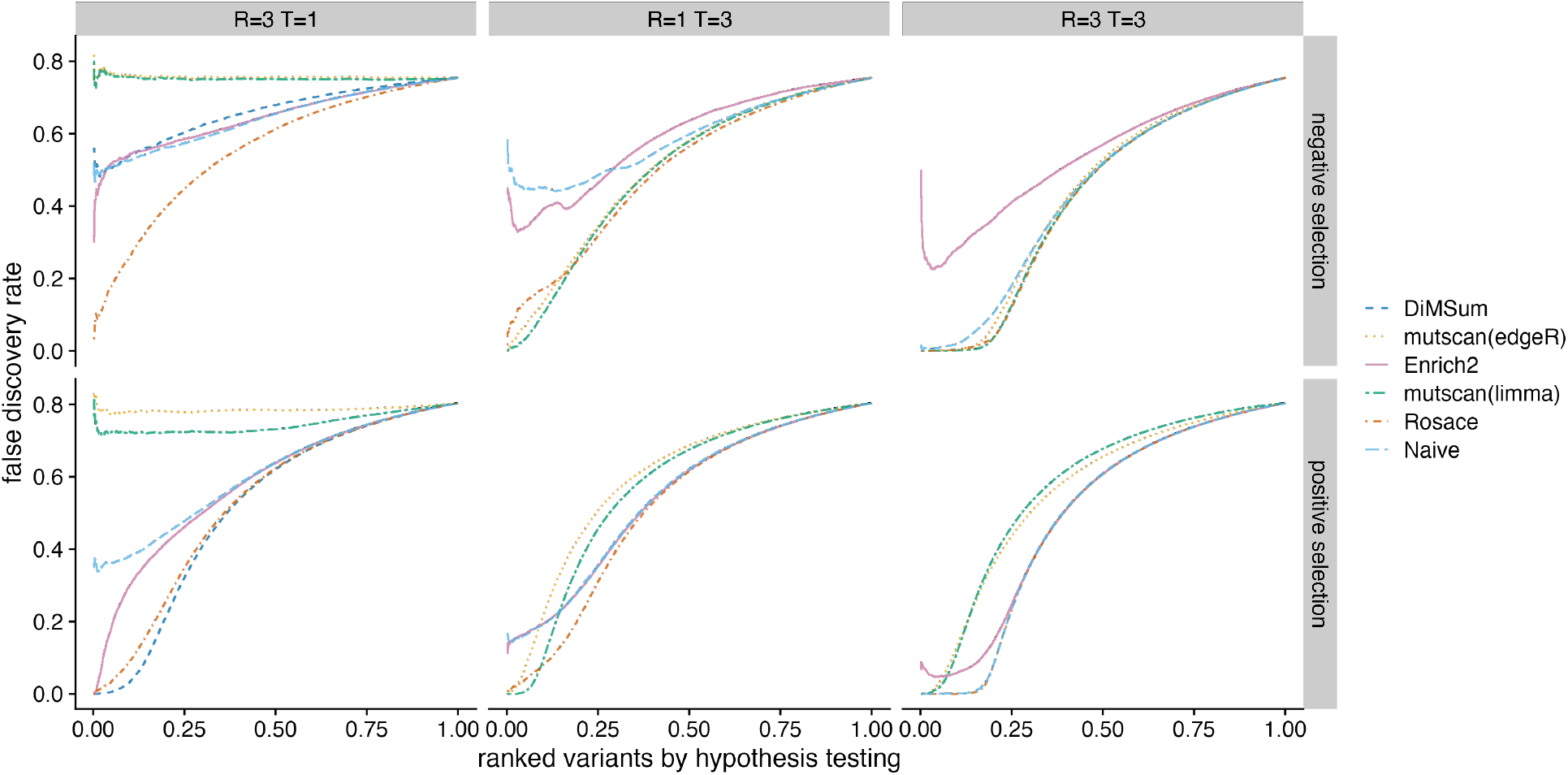
Benchmark of false discovery control on `Rosette` simulation. Variants are ranked by hypothesis testing (adjusted p-values or *lfsr*). The false discovery rate at each rank is computed as the proportion of neutral variants assuming all the variants till the rank cutoff are called significant. *R* is the number of replicates and *T* is the number of selection rounds. MET data is used for negative selection and OCT1 data for positive selection. Ideally, the line would be flat at 0 until the rank where all variants with true effects are discovered.

### 2.6 Testing Rosace Power with Rosette Simulation

Next, we investigate the sensitivity of benchmarking methods at different FDR or *lfsr* cutoff. It is important to keep in mind that `Rosace` uses raw *lfsr* from the sampling result while all other methods use the Benjamini-Hochberg Procedure to control the false discovery rate. As a result, the cutoff for `Rosace` is on a different scale.

`Rosace` is the only method that displays high sensitivity in all conditions with a low false discovery rate. In the case of one selection round and three replicates (*T* = 1 and *R* = 3), *edgeR* and *limma-voom* do not have the power to detect any significant variants with the FDR threshold at 0.1. The same scenario occurs with *DiMSum* at negative selection and the naive method at *T* = 3 and *R* = 1 (Fig. 6). The naive method in general has very low power, while *Enrich2* has a very inflated FDR.

**Figure 6.**
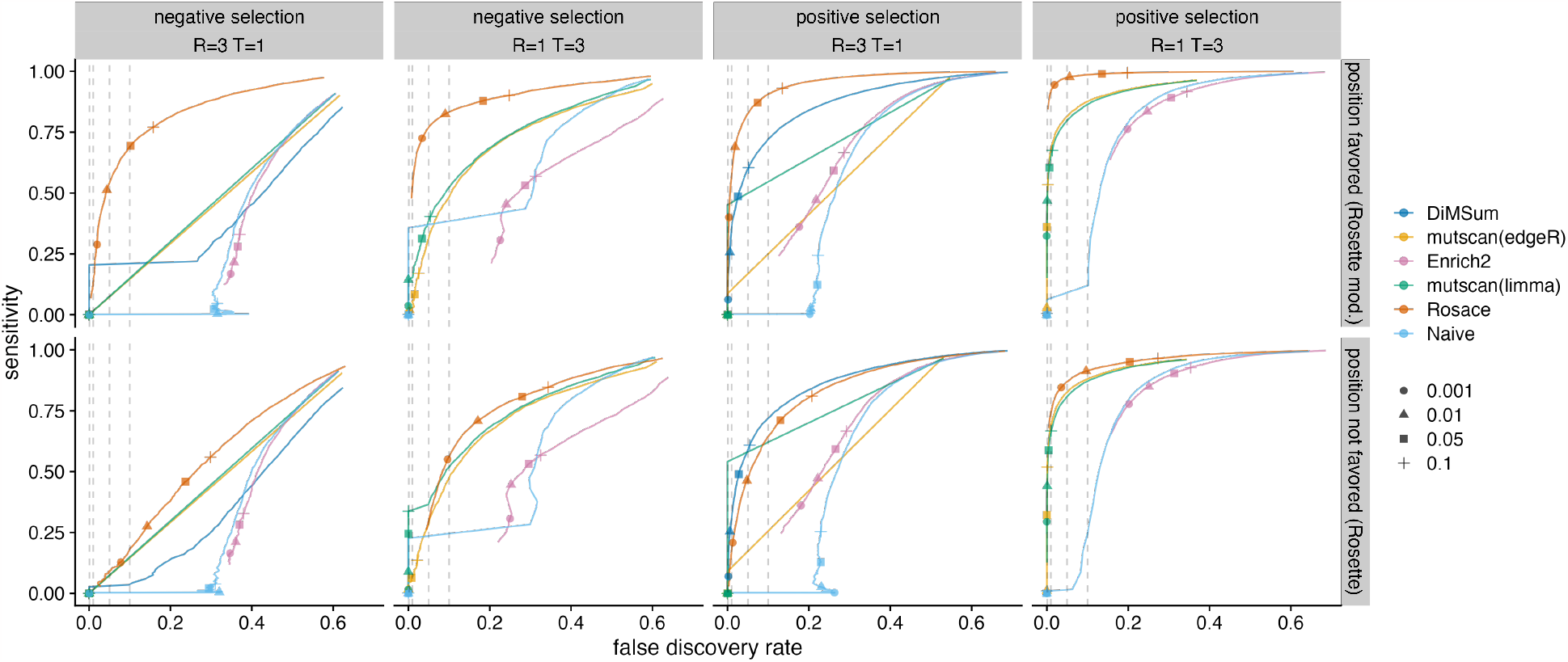
Benchmark of sensitivity versus FDR. The upper row is simulated from a modified version of `Rosette` simulation to favor position-informed models. The bottom row is the results from standard `Rosette`. Circles, triangles, squares, and crosses represent LOF variant selection at adjusted p-values or *lfsr* of 0.001, 0.01, 0.05, and 0.10, respectively. Variants with the opposite sign of selection are then excluded. Ideally, for all methods besides `Rosace`, each symbol would lie directly above the corresponding symbol on the x-axis indicating true FDR. For `Rosace`, *lfsr* has no direct translation to FDR so the cutoff represented by the shape is theoretically on a different scale.

We benchmark `Rosace` on both `Rosette` simulations, which inherently violate the position assumption, and a modified version of `Rosette` that favors the position-informed model. We show that model misspecification does increase the false discovery rate of `Rosace`, but `Rosace` is robust enough to outperform all other methods (except for *DiMSum* with *T* = 1 and *R* = 3 and positive selection) even when the position assumption is strongly violated (Fig. 6).

## 3 Discussion

We present `Rosace`, a Bayesian framework for analyzing growth-based deep mutational scanning data. In addition, we develop `Rosette`, a simulation framework that recapitulates the properties of actual DMS experiments, but relies on an orthogonal data generation process from `Rosace`. From both simulation and real data analysis, we show that `Rosace` has better FDR control and higher sensitivity compared to existing methods and that it provides reliable estimates for downstream analyses.

One of `Rosace`’s contributions is accounting for positional information in DMS analysis. The model assumes the prior information that variants on the same position have similar functional effects, resulting in higher sensitivity and better FDR. Furthermore, `Rosace` is also capable of incorporating other types of prior information on the similarity of variants.

The development of DMS simulation frameworks such as `Rosette` can also drive experimental design. For example, to select the best number of time points and replicates with regard to the trade-off between statistical robustness and costs of the experiment, an experimentalist can conduct a pilot experiment and use its data to infer summary statistics through `Rosette`. `Rosette` will then generate simulations close to a real experiment. Experimentalists can find the optimal tool for data analysis given an experimental design by applying candidate tools to the simulation data. Similarly, given a data analysis framework, experimentalists can choose from multiple experiment designs by using `Rosace` to simulate all those experiments and observe if any designs have enough power to detect most of the LOF or GOF variants with low false discovery rate.

This paper only applies our tool to growth screens, one of several functional phenotyping methods possible by DMS techniques. Other methods include fluorescently activated cell sorting (FACS-seq) - a branch of literature uses binned FACS-seq screens to sort the variant libraries based on protein phenotypes and therefore captures the distributional change of molecular properties [9, 4, 21]. Although of different design, FACS-seq-based screens can also be analyzed using a framework similar to `Rosace`. Building such frameworks incorporating prior information for experiments beyond growth screens enables the community to exploit a wider range of experimental data.

As the function of a protein is rarely one-dimensional, one can measure multiple phenotypes of a variant in a set of experiments. For example, the OCT1 data mentioned earlier [4] measures both the transporter surface expression from a FACS-seq screen and drug cytotoxicity with a growth screen. Multi-phenotype DMS experiments also call for analysis frameworks to accommodate multidimensional outcomes by modeling the interaction or the correlation of phenotypes of each variant. One successful attempt models the causal biophysical mechanism of protein folding and binding [22], and there are many more protein properties other than those two. The `Rosace` framework can also be adapted to such an extension. Hypothetically, if the MET inhibitor experiment treats cells in multiple chemicals separately before sequencing and the researcher believes the variants that are treated by the same chemical or genetic perturbations should have similar outcomes, such beliefs can be readily incorporated into the `Rosace` framework just as the belief that variants at the same location yield similar outcomes is built into it. Subsequently, `Rosace` allows for more prior knowledge about the relationship between perturbations and phenotypes, for example, by encoding protein and chemical properties, which leads to more general models. As a result, frameworks such as `Rosace` will help learn how across multi-dimensional relationships between genotypes and how genotypes give rise to emergent properties of biological systems. Then, we can better determine the effect of mutations and ultimately perhaps develop mechanistic or probabilistic models for how mutations drive proteins in evolution, how they lead to malfunction and diseases, and how to better engineer new proteins.

## 4 Methods

### 4.1 Pipeline: Raw Read to Sequencing Count

To facilitate the broader adoption of the `Rosace` framework for DMS experiments, we have developed a sequencing pipeline for short-read-based experiments using Snakemake which we dub Dumpling [23]. This pipeline handles directly sequenced single-variant libraries containing synonymous, missense, nonsense, and multi-length indel mutations, going from raw reads to final scores and quality control metrics. Raw sequencing data in the form of fastq files is first obtained as demultiplexed paired-end files. The user then defines the experimental architecture using a csv file defining the conditions, replicates, and time points corresponding to each file, which is parsed along with a configuration file. The reads are processed for quality and contaminants using BBDuk, and then the paired reads are error-corrected using BBMerge. The cleaned reads are then mapped onto the reference sequence using BBMap [24]. Variants in the resulting SAM file are called and counted using the AnalyzeSaturationMutagenesis tool in GATK v4 [25]. This tool provides a direct count of the number of times each distinct genotype is detected in an experiment. We generate various QC metrics throughout the process and combine them using MultiQC for an easy-to-read final overview [26].

Due to the degeneracy of indel alignments, the genotyping of codon-level deletions sometimes does not hew to the reading frame due to leftwise alignment. Additionally, due to errors in oligo synthesis, assembly, during in vivo passaging or during sequencing, some genotypes that were not designed as part of the library may be introduced. A fundamental assumption of DMS is the independence of individual variants, and so to reduce noise and eliminate error our pipeline removes those that were not part of our planned design before analysis, as well as renames variants to be consistent at the amino acid level, before exporting the variant counts in a format for `Rosace`.

We generate various QC metrics throughout the process and combine them using MultiQC for an easy-to-read final overview [26].

### 4.2 Pre-processing of Sequencing Count

In a growth DMS screen with *V* variants, we define *v* to be the variant index. A function *p*(*v*) maps the variant *v* to its position label. *T* indicates the number of selection rounds and index *t* is an integer ranging from 0 to *T*. A total of *R* replicates are measured, with *r* as the replicate index. We denote *c*_*v,t,r*_ the raw sequencing count of cells with variant *v* at time point *t* in replicate *r*.

In addition, “mutant” refers to substitution with one of the 20 amino acids, insertion of an amino acid, or deletion. Thus, a variant is uniquely identified by its mutant and the position where the mutant occurs (*p*(*v*)).

The default pre-processing pipeline of `Rosace` includes four steps: variant filtering, count imputation, count normalization, and replicate integration. First, variants with more than 50% of missing count data are filtered out in each replicate. Then, variants with a few missing data (less than 50%) are imputed using either the K-nearest neighbor averaging (*K* = 10) or filled with 0. Next, imputed raw counts are log-transformed with added pseudo-count 1/2 and normalized by the wild-type cells or the sum of sequencing counts for synonymous mutations. This step, which is proposed by *Enrich2*, allows for the computed functional score of wild-type cells to be approximately 0. Additionally, the counts for each variant before selection are aligned to be 0 for simple prior specification of the intercept.

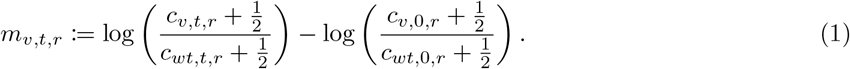

Previous papers suggest the usage of other methods such as total-count normalization when the wild-type is incorrectly estimated or subject to high levels of error [10, 12]. We include this in `Rosace` as an option. Finally, replicates in the same experiment are joined together for the input of the hierarchical model. If a variant is dropped out in some but not all replicates, `Rosace` imputes the missing replicate data with the mean of the other replicates.

### 4.3 Rosace: Hierarchical Model and Functional Score Inference

`Rosace` assumes that the aligned counts are generated by the following time-dependent linear function. Let *β*_*v*_ be the defined functional score or slope, *b*_*v*_ be the intercept, and *ϵ*_*g*(*v*)_ be the error term. The core of `Rosace` is a linear regression:

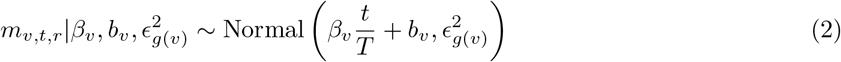

where *g*(*v*) maps the variant *v* to its mean group - the grouping method will be explained below.

*p*(*v*) is the function that maps a variant *v* to its amino acid position. If the information of variants’ mutation types is given, `Rosace` will group all the synonymous mutations together as a control position. In addition, we regroup positions with fewer than 10 variants together to avoid having too few variants in a position. For example, if the DMS screen has fewer than 10 mutants per position, adjacent positions will be grouped to form one position label.

We assume that the variants at the same position are more likely to share similar functional effects. Thus, we build the layer above *β*_*v*_ using position-level parameters *ϕ*_*p*(*v*)_ and *σ*_*p*(*v*)_.

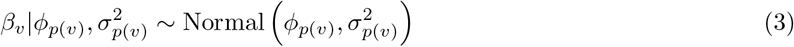

The mean and precision parameters are given a weakly-informative normal prior and variance parameters are given weakly-informative inverse-gamma distribution.

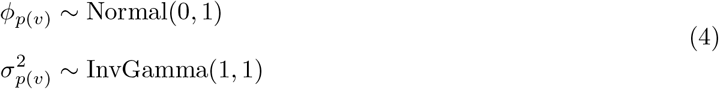

We further cluster the variant into mean groups of 25 based on its value of mean count across time points and replicates. The mapping between the variant and its mean group is denoted as *g*(*v*). Thus, we model the mean-variance relationship by assuming variants with a lower mean are expected to have higher error terms in the linear regression and vice versa.

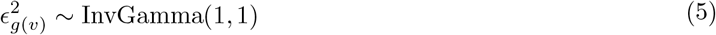

Stan [27] is used in `Rosace` for Bayesian inference over our model. We use the default inference method, the No-U-Turn sampler (NUTS), a variant of the Hamiltonian Monte Carlo (HMC) algorithm. Compared to other widely-used Monte Carlo samplers, for example, the Metropolis-Hastings algorithm, HMC has reduced correlation between successive samples, resulting in fewer samples reaching a similar level of accuracy [28]. NUTS further improves HMC by automatically determining the number of steps in each iteration of HMC sampling to more efficiently sample from the posterior [29].

The lower bound of the number of mutants per position index |{*v*|*p*(*v*) = *i*}| (10) and the size of the variant’s mean group *g*_*p*_ (25) can be changed.

### 4.4 Rosette: the OCT1 and MET Datasets

We use the following datasets as input of the `Rosette` simulation: the OCT1 dataset by Yee, Macdonald, and Mitrovic *et al*. [4] as an example of positive selection and the MET dataset by Estevam *et al*. [5] as an example of negative selection. Specifically, we use replicate 2 of the cytotoxicity selection screen in the OCT1 dataset for both score distribution and raw count dispersion. For the MET dataset, we select the experiment with IL-3 withdrawal under wildtype genetic background (without exon 14 skipping). Raw counts are extracted from replicate 1 but the scores are calculated from all three replicates because of the frequent dropouts at the initial time point.

The sequencing reads and the resulting sequencing counts are processed in the default pipeline described in the previous method sections. Scores are then computed using simple linear regression (the naïve method). The naïve method is used as the `Rosette` input because we are trying to learn the global distribution of the scores instead of identifying individual variants and, while uncalibrated, naïve estimates are unbiased.

### 4.5 Rosette: Summary Statistics from Real Data

Summary statistics inferred by `Rosette` can be categorized into two types: one for the dispersion of sequencing counts and the other for the dispersion of score distribution.

First, we estimate dispersion *η* in the sequencing count. We assume the sequencing count at time point 0 reflects the true variant library before selection. Since the functional scores of synonymous variants are approximately 0, the proportion of synonymous mutations in the population should approximately be the same after selection. Let the set of indices of synonymous mutations be **v**_*s*_ = {*v*_*s*1_, *v*_*s*2_, … }. The count of each synonymous mutation at time point *t* is 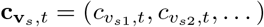. The model we use to fit *η* is thus

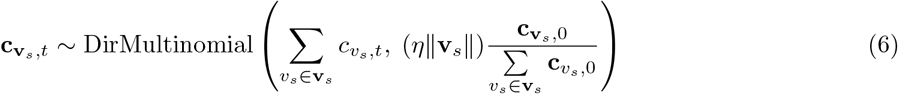

from which we find the maximum likelihood estimation 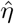.

Dispersion of the initial variant library *η*_0_ is estimated similarly by fitting a Dirichlet-Multinomial distribution on the sequencing counts of the initial time point assuming that in an ideal experiment, the proportion of each variant in the library should be the same. Similar to above, the indices of all mutations are **v** = {1, 2, …, *V* }, and the count of each mutation at time point 0 is **c**_**v**,0_ = (*c*_1,*t*_, *c*_2,*t*_, …, *c*_*V,t*_). From the following model

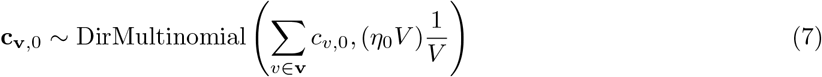

we can again find the maximum likelihood of the variant library dispersion 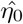. Notice that 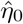 is usually much smaller than 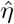 (i.e. more overdispersed) because 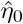 contains both the dispersion of the variant library as well as the sequencing step.

To characterize the distribution of functional scores, we first cluster mutants into groups, as mutants often have different properties and exert different influences on protein function. We calculate the empirical Jensen-Shannon divergence (JSD) to measure the distance between two mutants, using bins of 0.1 to find the empirical probability density function. Ideally, a clustering scheme should produce a grouping that reflects the inherent properties of an amino acid that are independent of position. Thus, we are more concerned with the general shape of the distribution than the similarity between paired observations. It leads to our preference for JSD over Euclidean distance as the clustering metric. To cluster mutants into four mutant groups *g*_*m*_ = {1, 2, 3, 4}, we use hierarchical clustering (“hclust” function with *complete linkage* method in R), and we record the proportions 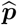 to simulate any number of mutants in the simulation (the number of mutant groups can also be changed). The underlying assumption is that mutants in each mutant group are very similar and can be treated as interchangeable. We define *f*_1_(*v*) as the function that maps a variant to its corresponding mutant group *g*_*m*_.

Then we cluster the variants into different variant groups. In the case of our examples, the shape is not unimodal but bimodal. The OCT1 screen has a LOF mode on the right (positive selection) and the MET screen has a LOF mode on the left (negative selection). While it is possible to observe both GOF and LOF variants, we observed in our datasets that GOF variants are so rare that they do not constitute a mode on the mixed distribution, resulting in a bimodal distribution. To cluster the non-synonymous variants into groups *g*_*v*_, we use the Gaussian Mixture model with two mixtures for our examples to decide the cutoff of the groups, and we fit the Gaussian distribution for each variant group again to learn the parameters of the distribution. The synonymous variants have their own group labeled as control. Let *f*_2_(*v*) denote the function that maps a variant to its corresponding variant group *g*_*v*_. The result of the simulation shows that even the synonymous mutations with scores close to 0 can have large negative effects due to random dropout. Thus, we later set the effect of the control and the neutral group to be constant 0 and still observe a similar distribution as seen in the real data. For each variant, we have one of the models below, depending on whether the variant results in LOF or has no effects:

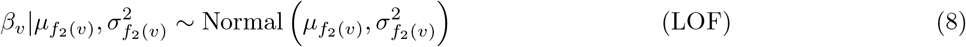

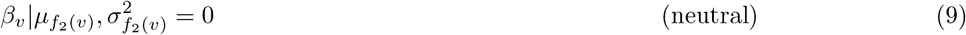

We use 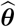 to denote the collection of estimated distributional parameters for all variant groups. Finally, we define the number of variants in each variant group at each position

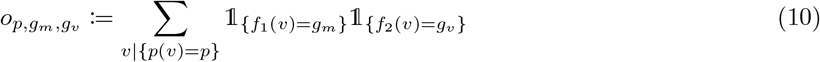

For each position *p*, we can thus find the count of variants belonging to any mutant-variant group 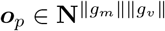. Treating each position as an observation, we fit a Dirichlet distribution to characterize the distribution of variant group identities among mutants at any position:

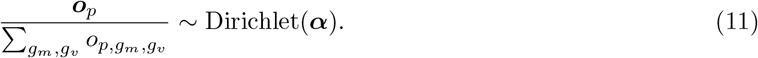

The final summary statistics are 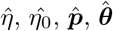, and 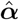. We also need *T*, the number of selection rounds, to map *β*_*v*_ into the latent functional parameter *μ*_*v*_ in growth screens.

### 4.6 Rosette: Data Generative Model

We simulate as the real experiment the same number of mutants *M*, the number of positions *P*, and the number of variants *V* (*M × P*). The important hyperparameters that need to be specified are the average number of reads per variant *D* (100, also referred to as the sequencing depth), initial cell population count *P*_0_ (200*V*), and wild-type doubling rate *δ* between time points (−2 or 2). One also needs to specify the number of replicates *R* and selection rounds *T*.

The simulation largely consists of two major steps: 1) generating latent growth rates *μ*_*v*_ and 2) generating cell counts *N*_*v,t,r*_ and sequencing counts *c*_*v,t,r*_.

In step 1, the mutant group and variant group labeling of each variant is first generated. Specifically, we assign a mutant to the mutant group *g*_*m*_ by the proportion 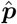 and then assign a variant to the variant group *g*_*v*_ by drawing o_*p*_ from Dirichlet distribution with parameter 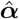(Equation 10). Using 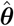, we randomly generate *β*_*v*_ for each variant based on its *g*_*v*_ (Equation 8). The mapping between *β*_*v*_ and *μ*_*v*_ requires an understanding of the generative model, so it will be defined after we present the cell growth model.

In step 2, the starting cell population *N*_*v,r*,0_ is drawn from a Dirichlet-Multinomial distribution using 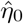 and we assume that replicates are biological replicates:

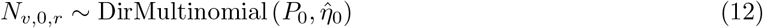

where *P*_0_ is the total cell population. The cells are growing exponentially and we determine the cell count by a Poisson distribution

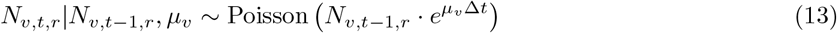

where Δ*t* is the pseudo-passing time. It differs from index *t* and will be defined in the next paragraph. Similar to how we define **c**_**v**,*t,r*_, we define the true cell count of each variant at time point *t* and replicate *r* to be **N**_**v**,*t,r*_ = (*N*_1,*t,r*_, …, *N*_*V,t,r*_). The sequencing count for each variant is

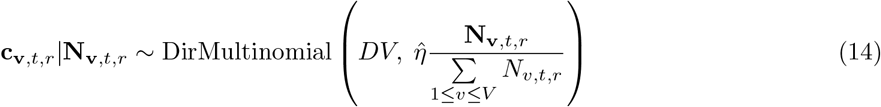

where *D* is the sequencing depth per variant. Empirically, we can set input 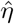 and 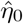 slightly higher than the estimated summary statistics. This is because the estimated values encompass all the noises in the experiment, while the true values only represent the noise from the sequencing step.

To find the mapping between *β*_*v*_ and *μ*_*v*_, we define *δ* to be the wild-type doubling rate and naturally compute 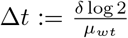, the pseudo-passing time in each round. Then we can compute the expectation of *β*_*v*_ with the linear regression model. For simplicity, we omit the replicate index *r* and assume *r* is fixed in the next set of equations.

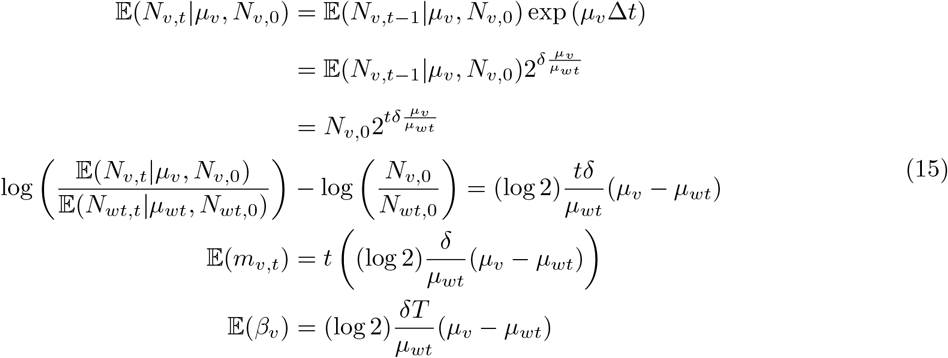

The final mapping between simulated *β*_*v*_ and *μ*_*v*_ is then described in the following

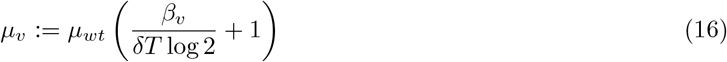

with *μ*_*wt*_ set to be sgn(*δ*).

### 4.7 Modified Rosette that Favors Position-Informed Models

In the original, position-agnostic version of `Rosette`, a ∥*g*_*m*_∥∥*g*_*v*_∥-dimensional vector is drawn from the same Dirichlet distribution for each position. The vector can be regarded as a quota for each mutant-variant group. Variants at each position are assigned their mutant-variant group according to the quota. As a result, at one position, variants from all variant groups (neutral, LOF, and GOF) would exist, and this violates the assumption in `Rosace` that variants at one position would have similar functional effects (strong LOF and GOF variants are very unlikely to be at the same position). To show that `Rosace` could indeed take advantage of the position information when it exists in the data, we create a modified version of `Rosette` where variants at one position could only belong to one variant group. Specifically, a position can have either neutral, LOF, or GOF variants, but not a mixture among any variant groups.

### 4.8 Benchmarking

The naïve method (simple linear regression) is conducted by the “lm” function in R on processed data. For each variant, normalized counts are regressed against time. Raw two-sided p-values are computed from t-statistics given by the “lm” function. It is then corrected using the Benjamini-Hochberg Procedure to adjust the p-values.

For *Enrich2*, we use the built-in variant filtering and wild-type (“wt”) normalization. All analyses use a random-effect model as presented in the paper. When there is more than one selection round, we use weighted linear regression. Otherwise, a simple ratio test is performed. The resulting p-values are adjusted using the Benjamini-Hochberg Procedure.

*DiMSum* requires the variant labeling to be DNA sequences. As a result, we have to generate dummy sequences. It is applied to all simulations with one selection round with the default settings. The z-statistics are computed using the variant’s mean estimate over the estimated standard deviation and the adjusted p-value is computed from the z-score with Benjamini-Hochberg Procedure.

*mutscan* is an end-to-end pipeline that requires the input to be sequencing reads. Conversely, `Rosette` only generates sequencing counts, which can be calculated from sequencing reads but cannot be used to recover sequencing reads. To facilitate benchmarking, we use a *SummarizedExperiment* object to feed the `Rosette` output to their function “calculateRelativeFC”, which does take sequencing counts as input. We benchmark both *edgeR* and *limma-voom* with default normalization and hyperparameters as provided in the function. We use the “logFC shrunk” and “FDR” columns in *edgeR* output and the “logFC” and “adj.P.Val” columns in *limma-voom* output.

We run `Rosace` with position information of variants and labeling of synonymous mutations. However, `Rosace` is a Bayesian framework so it does not compute FDR like the frequentist methods above. All `Rosace` power/FDR calculations are done under the Bayesian local false sign rate (*lfsr*) setting [20]. As a result, in the simulation, we present the rank-FDR curve and the FDR-Sensitivity curve as the metrics instead of setting an identical or different hard threshold on FDR and *lfsr*. In the real data benchmarking, both the FDR and *lfsr* thresholds are set to be 0.05.

## Availability of data and materials

`Rosace` is implemented as an R package and is distributed on GitHub (https://github.com/pimentellab/rosace). The package also includes functions for `Rosette` simulation. The integrated sequencing pipeline for short-read-based experiments is available on GitHub (https://github.com/odcambc/dumpling). Scripts and public datasets used to perform data analysis and generate plots for the paper are uploaded on GitHub as well (https://github.com/roserao/rosace-paper-script).

## Author Information

### Authors and Affiliations

Department of Computer Science, UCLA, Los Angeles, CA, USA Jingyou Rao, Harold Pimentel

Computational and Systems Biology Interdepartmental Program, UCLA, Los Angeles, CA, USA Ruiqi Xin

Department of Bioengineering and Therapeutic Sciences, UCSF, San Francisco, CA, USA

Christian Macdonald, Matthew Howard, Gabriella O. Estevam, Sook Wah Yee, James S. Fraser, Willow Coyote-Maestas

Tetrad Graduate Program, UCSF, San Francisco, CA, USA

Matthew Howard, Gabriella O. Estevam

Department of Pharmaceutical Chemistry, UCSF, San Francisco, CA, USA

Matthew Howard

Department of Mathematics, Baruch College, CUNY, New York, NY, USA

Mingsen Wang

Quantitative Biosciences Institute, UCSF, San Francisco, CA, USA

James S. Fraser, Willow Coyote-Maestas

Department of Computational Medicine, David Geffen School of Medicine, UCLA, Los Angeles, CA, USA Harold Pimentel

Department of Human Genetics, David Geffen School of Medicine, UCLA, Los Angeles, CA, USA Harold Pimentel

## Contributions

JR, CM, WCM, and HP jointly conceived the project. JR and HP developed the statistical model and the simulation framework. JR, MW, and RX wrote the software and its support. JR performed the data analysis and benchmarking. CM wrote the sequencing pipeline. SWY and CM performed the OCT1 experiment and GOE performed the MET experiment. JR and HP wrote the manuscript with input from MW, CM, WCM, MH, and JF. All authors read and approved the final manuscript.

## Supplementary Figures

**Fig. S1:**
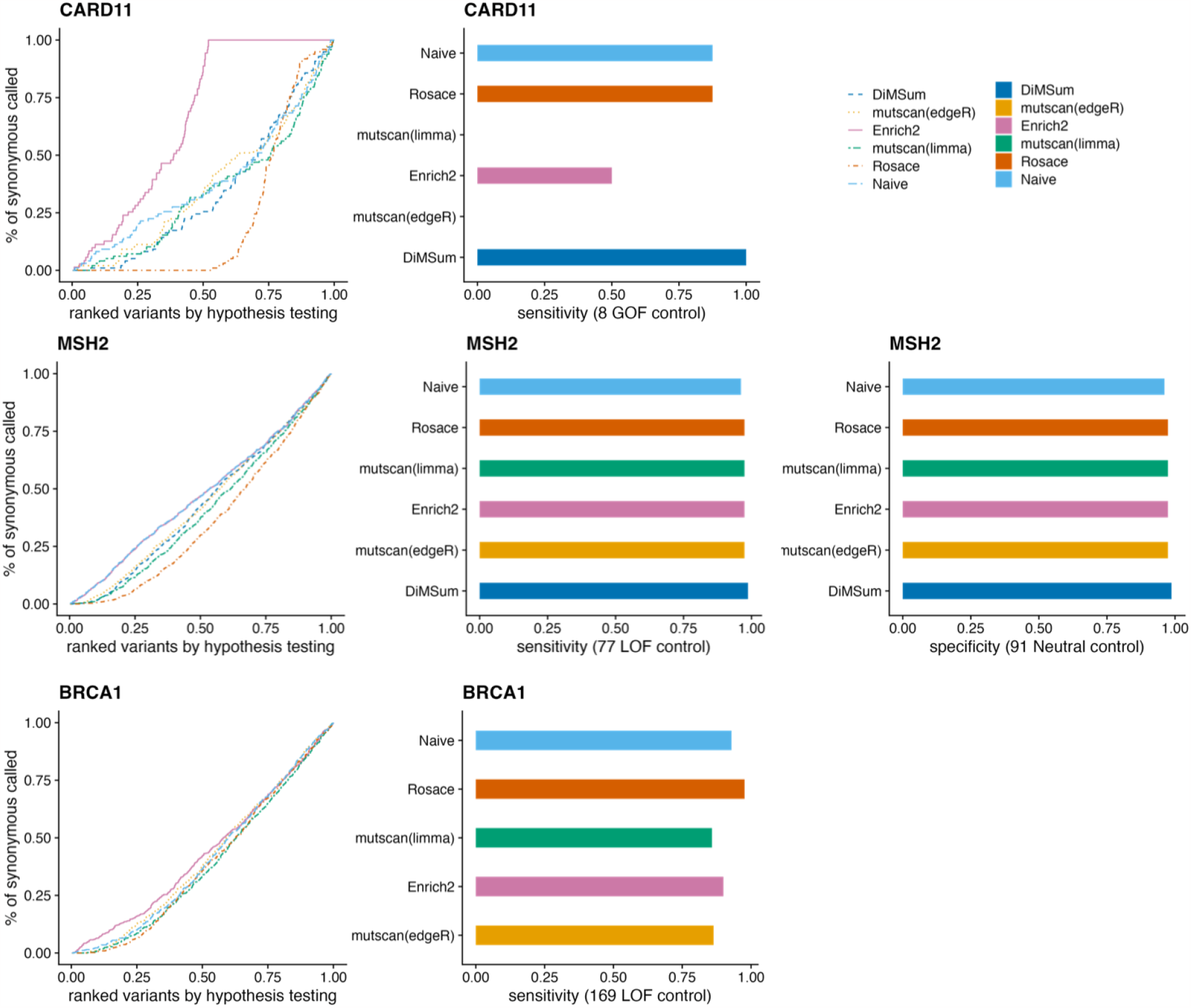
`Rosace` performance on CARD11 [8], MSH2 [6], and BRCA1 [7] datasets. Line plots in the leftmost panel are similar to Figure 3A in the main text, representing the false discovery rate of different methods. Bar plots show the sensitivity and specificity. The CARD11 validation set is experimentally validated in the paper with high confidence. The MSH2 and the BRCA1 validation sets are generated by the authors from ClinVar and other resources.

**Fig. S2:**
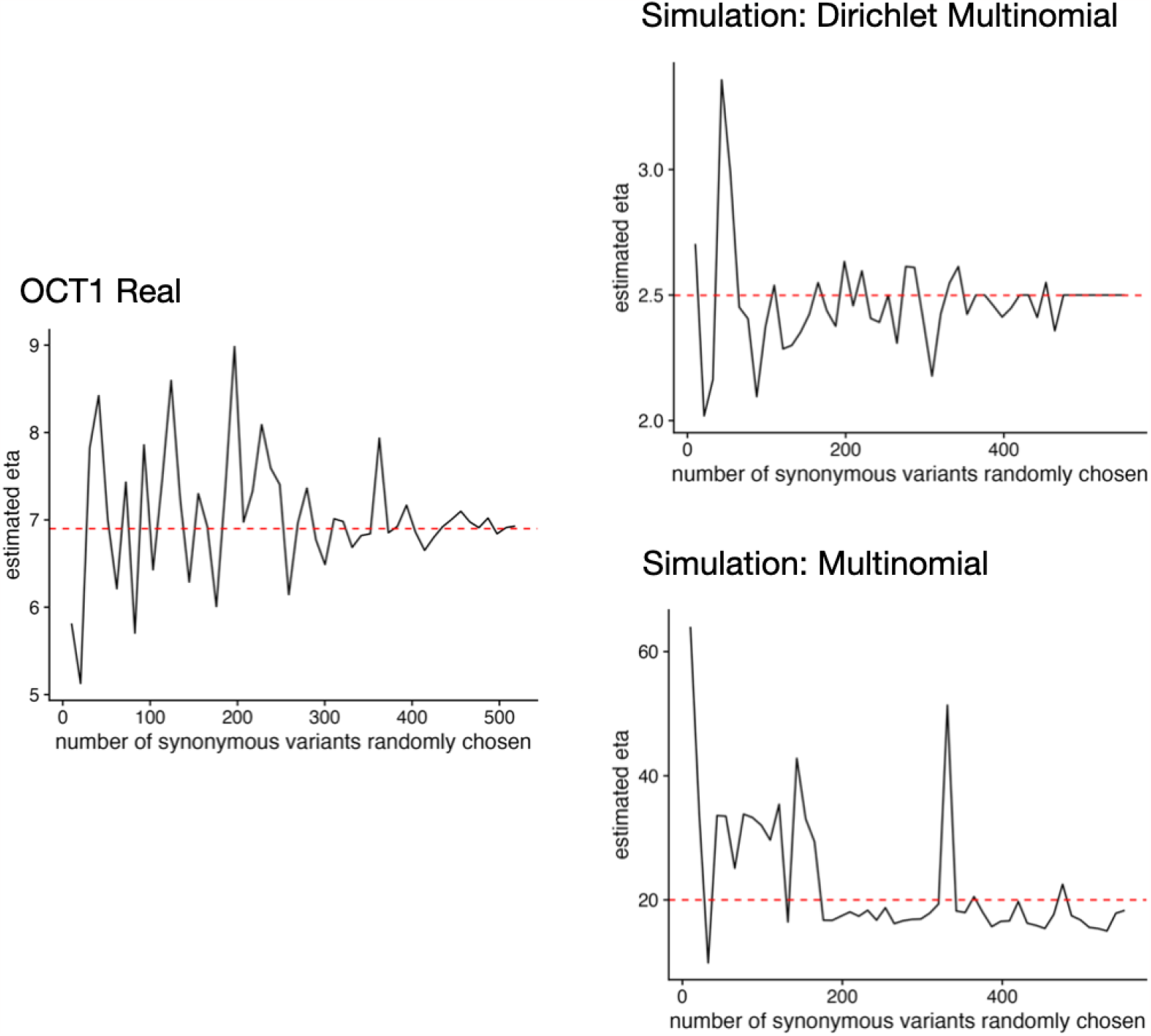
Dispersion of the sequencing counts. The multinomial distribution is not enough to characterize the large variability of proportion changes within the synonymous mutations, so instead we use the Dirichlet-Multinomial distribution to adjust for the overdispersion in the simulation. The y-axis is the estimated dispersion value and the x-axis is the number of synonymous randomly chosen as input to complete the estimation.

**Fig. S3:**
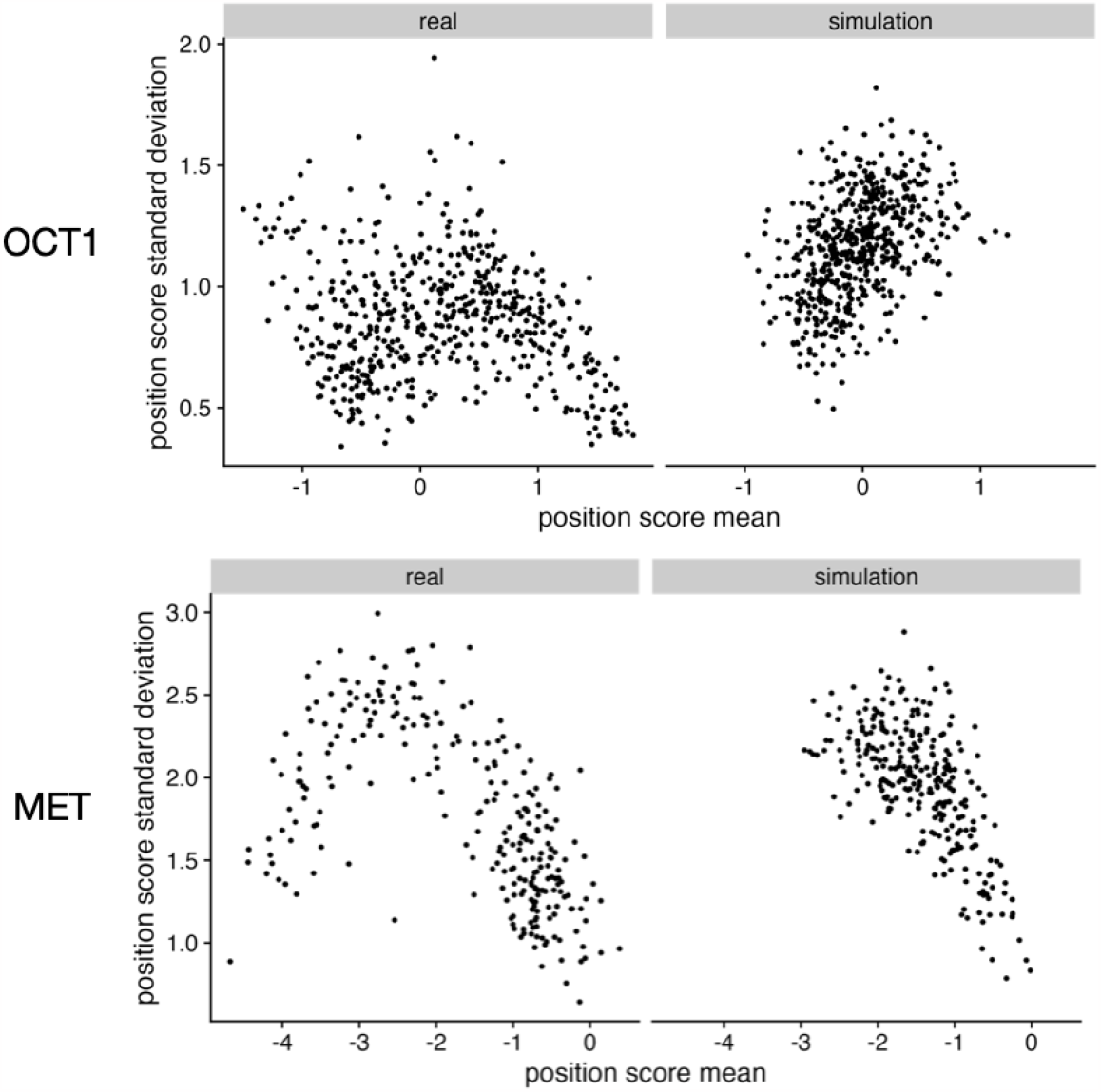
Comparison of position-level score profile between real and simulation data. In real data, the positions with extreme scores (very positive or negative scores) have reduced standard deviation, but this effect does not appear in the simulation.

**Fig. S4:**
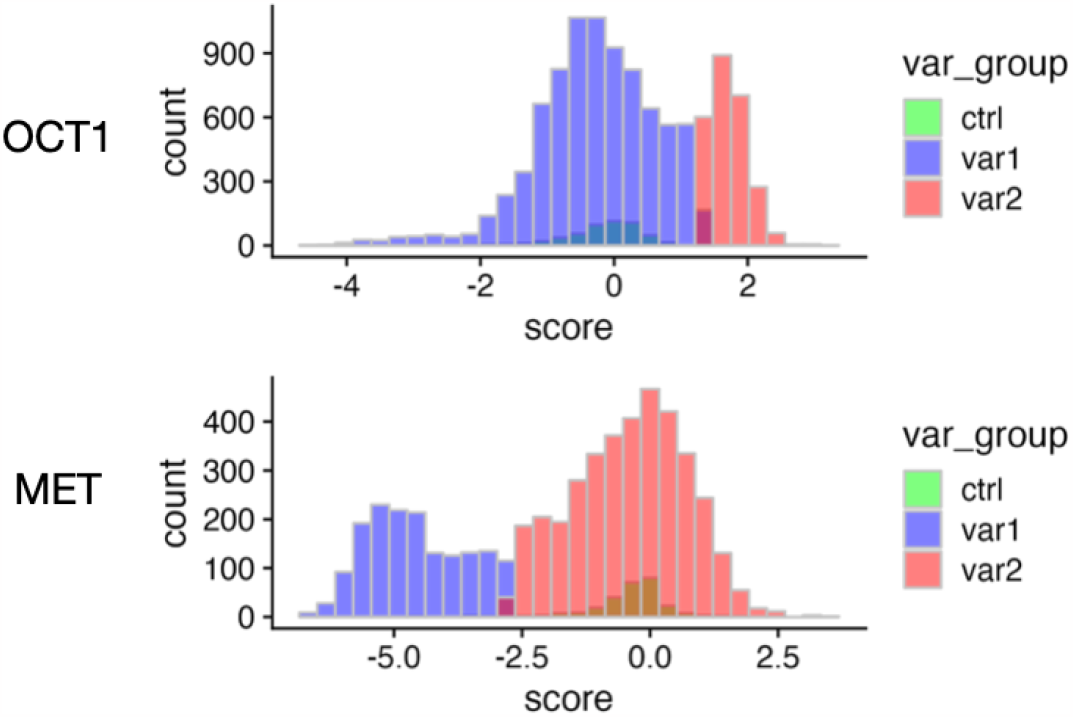
Both OCT1 and MET datasets have a clear separation of two modes in the functional score distribution (neutral and LOF, “var1” and “var2” in the legend). We treat OCT1 as a positive selection screen and MET as a negative selection screen.

**Fig. S5:**
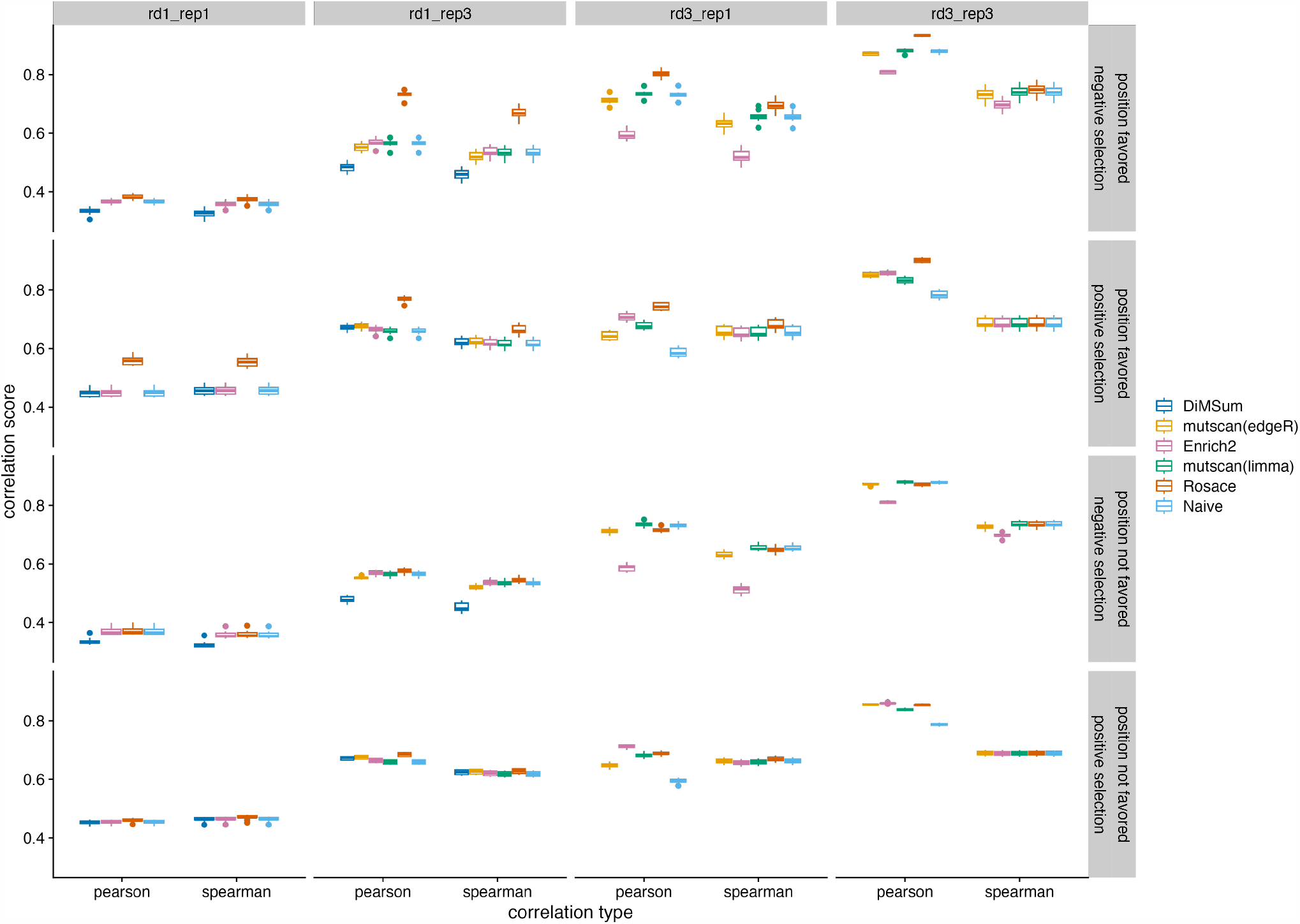
Comparison of correlation score on `Rosette` simulation. “rd*T*_rep*R*” stands for *T* selection rounds and *R* replicates.

**Fig. S6:**
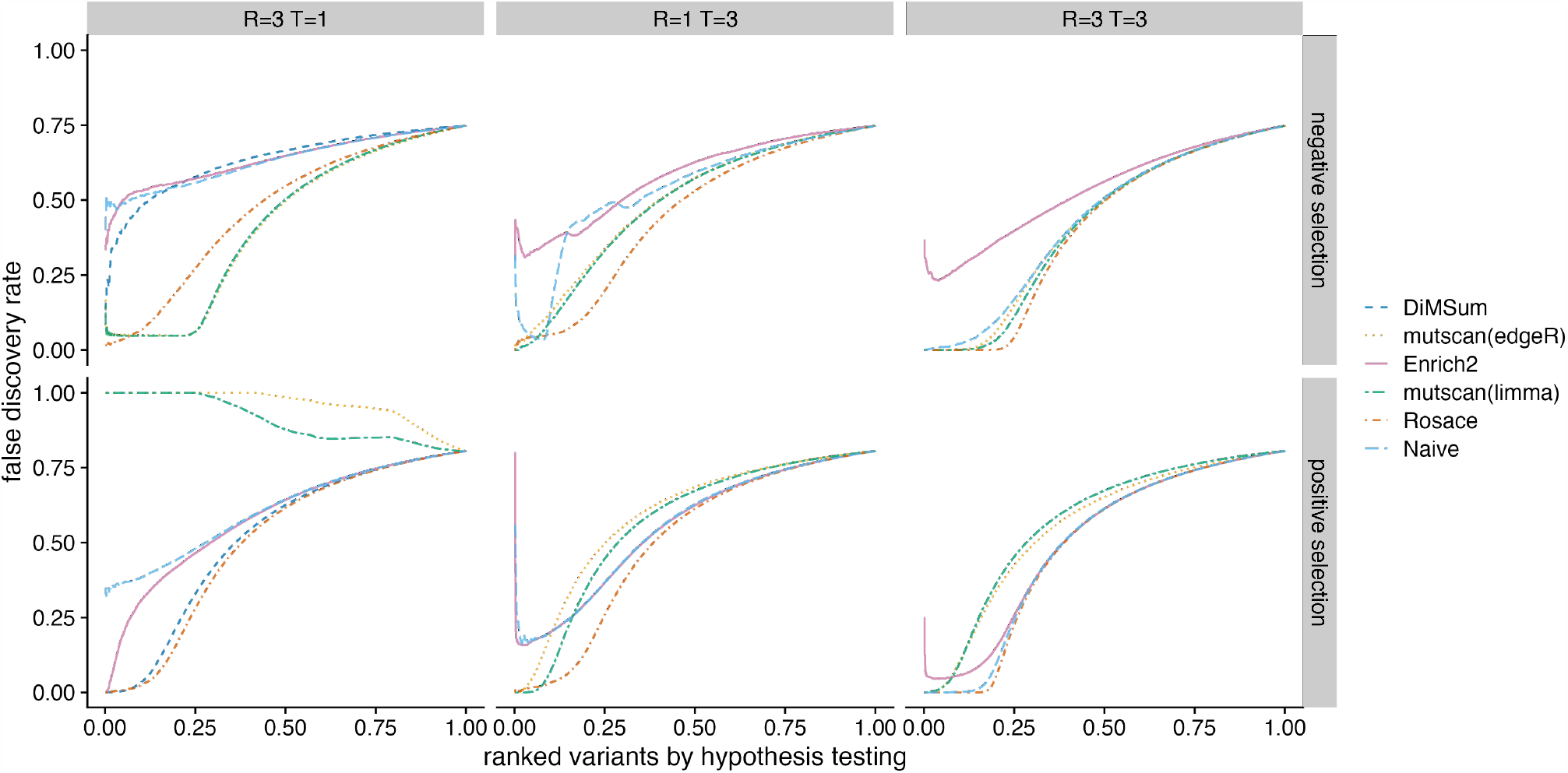
Benchmark of false discovery control on `Rosette` simulation under position-favored model. Similar to Figure 5 in the main text. *R* stands for the number of replicates and *T* stands for the number of selection rounds.

**Fig. S7:**
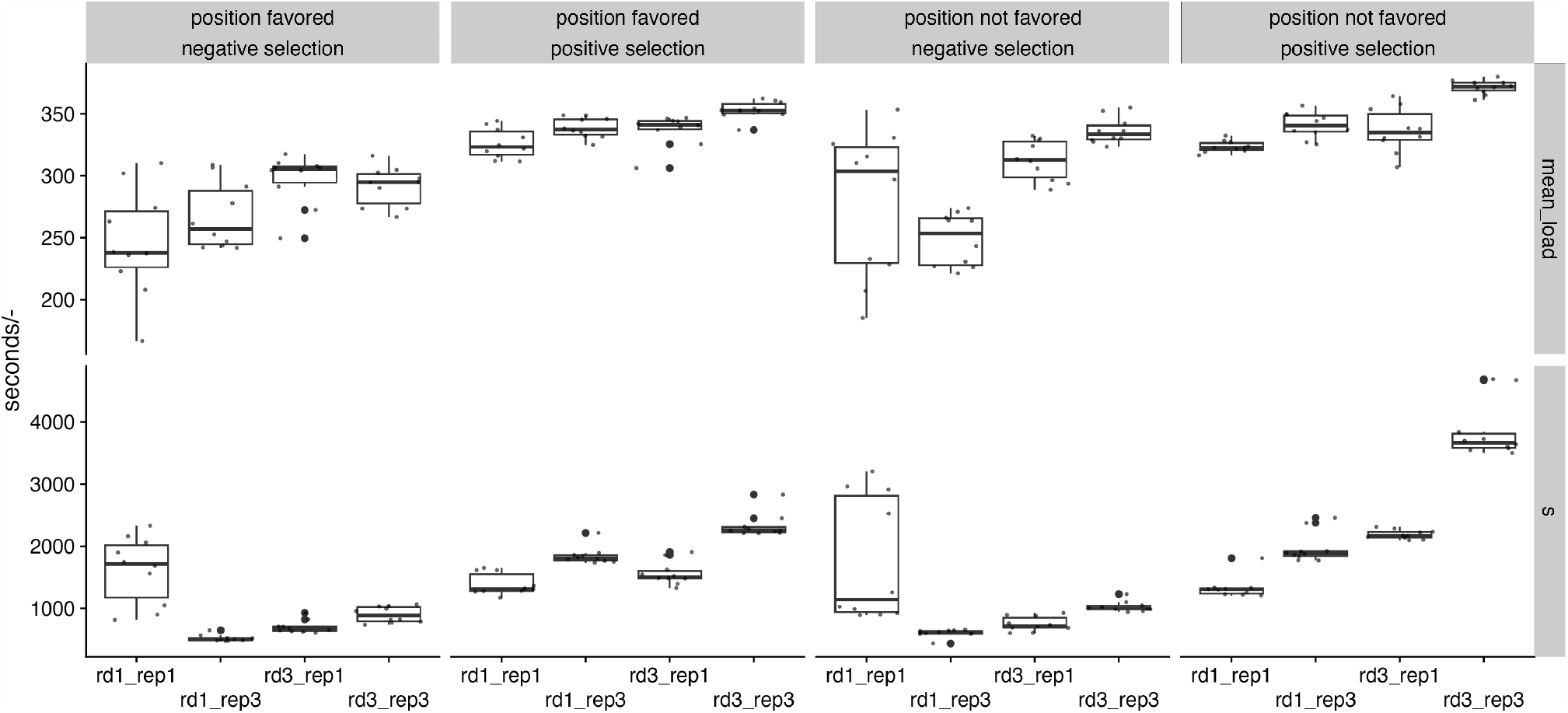
Running time in seconds and mean load (CPU usage over time, divided by the total running time) of `Rosace` program. “rd*T*_rep*R*” stands for *T* selection rounds and *R* replicates.

## Notes

https://github.com/pimentellab/rosace

### Competing Interest Statement

The authors have declared no competing interest.

